# Capturing secondary structure in coarse grained intrinsically disordered proteins with simulations driven by chemical shifts

**DOI:** 10.64898/2026.01.06.697719

**Authors:** Mina Cullen, Carmen Biancaniello, Katerina Taškova, Vedran Miletić, Alfonso De Simone, Davide Mercadante

**Affiliations:** School of Chemical Sciences, The University of Auckland, Auckland, New Zealand; Department of Pharmacy, University of Naples Federico II, Naples, Italy; Department of Computer Science, The University of Auckland, Auckland, New Zealand; Max Planck computing and data facility (MPCDF), Garching, Munich, Germany; Maurice Wilkins Centre for molecular biodiscovery, The University of Auckland, Auckland, New Zealand

## Abstract

A major challenge when investigating intrinsically disordered proteins (IDPs) pertains to understanding how secondary structure formation across otherwise disordered ensembles, relates to function.

While coarse-grained (CG) simulation methods are instrumental to describe ensemble-to-function relationships in IDPs, they remain largely incapable of sampling secondary structure.

We here present NapshiftC_α_, an artificial neural network capable of modelling NMR chemical shifts from C_α_-mapped CG protein conformations, leveraging its predictions to restrain simulations and yield experimentally-informed IDP ensembles that contain secondary structure.

We applied NapshiftC_α_ on a large set of proteins in isolation or condensed phase, and found that, when secondary structure is present, CS incorporation yields improved agreements with smFRET spectroscopy and resolves the over-reliance of CG models on near linear conformers.

By incorporating observables that are routinely collected for IDPs but never coupled to CG simulations, NapshiftC_α_ ultimately informs on both local and global conformational features of function-defining IDP ensembles.

## Introduction

The discovery of intrinsically disordered proteins (IDPs) has opened a new frontier of biology, where functional aspects of proteins are no longer described by well-defined structures or a set of few conformations, but by a myriad of fast interconverting conformers organised in functional sub-ensembles.^1,2^ The lack of stable secondary structure in proteins has concomitantly brought methodological challenges, as the extreme dynamics characterising IDPs makes most of the structural biology techniques unsuitable for their investigation. Studies targeting IDPs thus often revolve around the orthogonal combination of experimental techniques, such as nuclear magnetic resonance (NMR) and single-molecule Förster resonance energy (smFRET) spectroscopy: which are well-versed at informing on local conformations and transient inter- and intra- molecular contacts.^3–9^ In combination with those, computational methods have heavily contributed to the mechanistic understanding of IDPs^10^, often through a quantitative matching of computed and experimental observables.^5,11–16^

Some of the challenges pertaining to the computational investigation of IDPs relate to their extreme conformational dynamics and large size, which require big simulation boxes. Additionally, IDPs have a strong propensity to organise in bulk phases via liquid-liquid phase separation (LLPS), which recently have posed new challenges to structural biology.^17^

Simulations of IDPs have thus evolved towards the refinement of coarse-grained (CG) models capable of dealing with chains of larger size and systems of increasing complexity. Recent developments have seen the introduction of numerous CG models largely employing a C_α_-based mapping where each residue is modelled as a single bead of certain mass and radius. The force fields in support of this mapping have, as their focal point, the refinement of non-bonded parameters achieved through physics- or machine learning (ML)-based strategies, as well as a combination of both.^18–26^

By contrast, bonded interactions are represented using simplified harmonic potentials, whereas ensemble validation is largely reliant on comparisons with the radius of gyration (R_g_), as this is the most available experimental data for IDPs. Relevantly, recent efforts have sought to refine the dihedral potentials of these models to attempt the sampling of secondary structure.^27,28^

On the experimental side, NMR is routinely used for the study of IDPs and can offer a considerable source of data to validate CG force fields^29,30^, or to directly integrate its observables into simulations and guide the exploration of conformational ensembles incorporating secondary structure.^31^

Among the NMR observables available to elucidate the properties of IDPs, chemical shifts (CS) constitute a wealth of information and have been useful to inform cluster analyses on IDP ensembles.^32^ CS are strongly informative of local changes in the chemical environment of protein atoms and function as a sensitive probe of secondary structure, including transient structural elements which are a functional hallmark of IDPs.^29,33–36^ Importantly, the differentiability in cartesian space of models relating CS values to protein structures allowed to integrate these observables into structural biology and computational biophysics, through either the refinement of NMR experimental structures^37–39^ or by restraining all-atom MD simulations.^40,41^

For small systems, CS-restrained all-atom simulations have shown to accurately refine the energy landscape of IDPs.^42^ The feasibility of this approach, however, rapidly diminishes with system size, thereby requiring the adoption of simplified CG representations.

The challenge of integrating CS into CG simulations relates to the fundamental atomistic nature of CS, which is lost upon coarse graining. ML-based approaches especially those capable of learning complex relationships and multi-scale representations, are however uniquely suited to bridge the atomistic-to-CG gap. These models could thus be trained to reconstruct the missing atomistic context statistically, leveraging patterns learnt from all-atom data and learning how to map those onto a CG space.

We here leverage the ability of neural networks to be highly predictive of patterns learnt statistically, and present NapshiftC_α_: an artificial neural network (ANN) that can predict CS from C_α_-mapped protein chains. We then use the obtained predictive ability of NapshiftC_α_ to integrate CS into MD simulations of coarse grained IDPs, by predicting CS at each simulation step. The difference between experimental and computed CS is then used to restrain simulations by correcting the forces acting on the beads, as sampled by the underlying parametric force field adopted.

This restraining approach is shown to guide simulations of IDPs towards exploring conformational spaces complying with experimental CS and increasing the probability of sampling experimentally informed ensembles. As CS are strongly informative of secondary structure, NapshiftC_α_ drives a local rearrangement of beads leading to the formation of experimentally informed α-helices or β-strands, while the overall ensemble dimensions can be still largely dictated by the non-bonded parameters of the force field.

Ultimately, through an integrative structural biology approach, NapshiftC_α_ enriches the investigation of the ensemble-to-function paradigm in IDPs by sampling secondary structure in coarse graining and expanding ensembles’ functional interpretability as empowered by NMR.

## Results

### Napshit*C_α_* is an artificial neural network allowing the sampling of secondary structure in strongly coarse-grained representations of IDPs

We trained NapshiftC_α_ to predict CS for C_α_-mapped protein chains, as this topology reflects the modelling paradigm most commonly employed in simulations of IDPs. ^18,20,22,23,25,28,48–51^ For the training, we used a dataset of 3,236 protein structures resolved with NMR and equipped with associated CS data (see methods for details).

Given the close relationship between CS and dihedral angles, we analysed how exhaustively the NapshiftC_α_ dataset informs on the angular properties of the protein structures employed for the ANN training. We therefore first projected the dataset onto the Ramachandran plot and classified its regions according to the point density along φ and ψ dihedrals (Fig. 1a-b). As suggested previously^43^, within the Ramachandran plot we classify four populations: (i) dihedrals in strongly structured regions (either α-helix or β-strands) when point scatter density is ≥500, (ii) regions surrounding the core of structured regions when point scatter density is ≥100 and <500, (iii) allowed regions when point scatter density is ≥50 and <100 and (iv) generously allowed regions when point scatter density is ≥10 and <50. Beyond those are disallowed regions.

**Figure 1.**
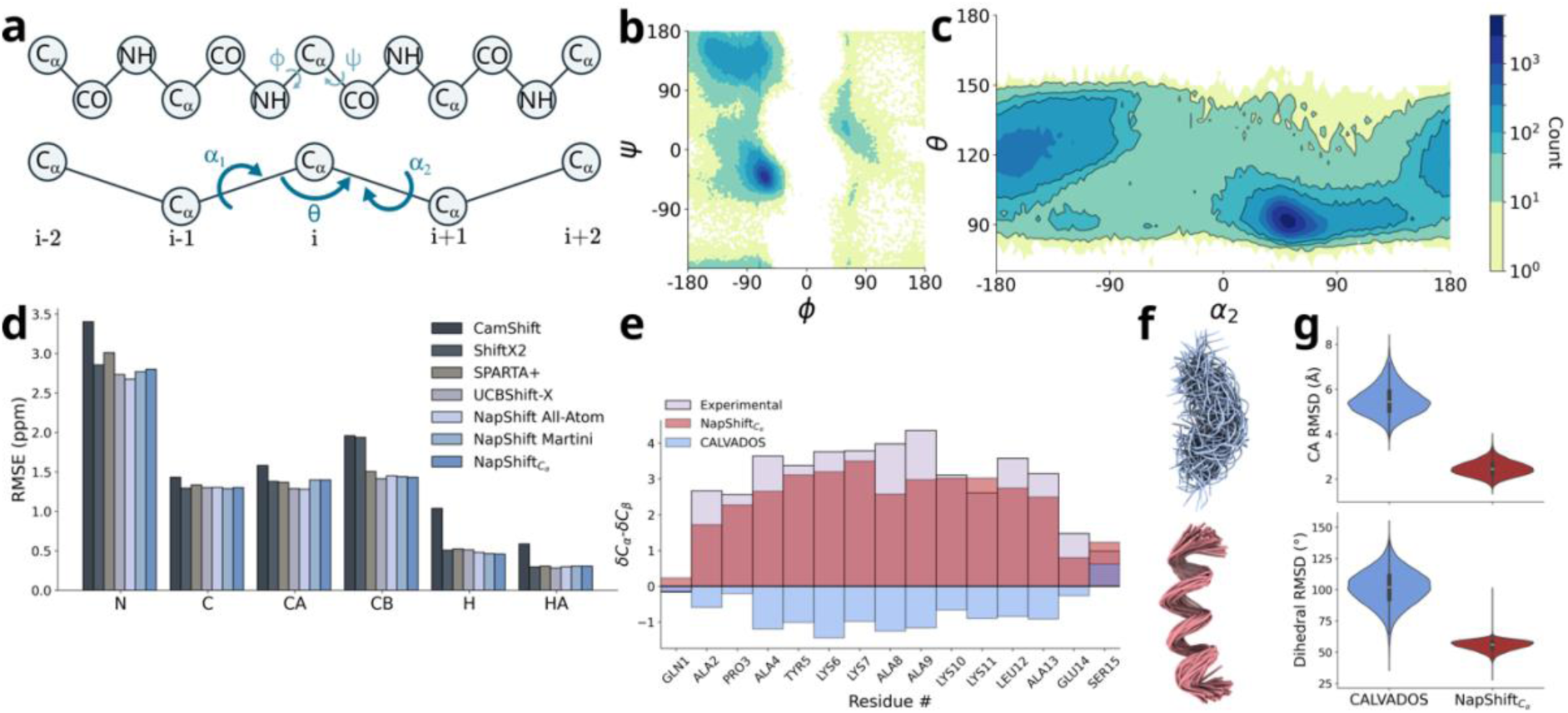
Training and performance of NapshiftC_α_. (a) Schematic representation of an all-atom (top panel) and coarse-grained C_α_-only (CG) representation of a protein chain, highlighting the angular features defining the chain’s conformational degrees of freedom. The φ and ψ angles dominate conformational propensities of all-atom chains and are proxied by α_1,2_ and θ in CG. (b-c) Ramachandran (b) and pseudoramachandran (c) plot of θ vs. α_2_ of the dataset used to train NapshiftC_α_, with entries coloured by count. The respective θ vs. α_1_ plot is reported in Figure S1. (d) Root mean square error (RMSE) of the predicted chemical shifts (CS) for each backbone atom, obtained from several CS predictors as shown in the legend. (e) Secondary chemical shifts for the α-helical peptide of the human platelet factor (PDB accession code 1DNG, BMRB accession code 4723). (f) representative ensembles simulated with unrestrained (Calvados only, blue) and restrained (NapshiftC_α_, red) simulations. (g) Cartesian (top panel) and dihedral (bottom panel) root mean square deviation (RMSD) of the peptide shown in (f) simulated with either unrestrained (blue) and restrained (red) simulations.

We then projected this angular space into the two main angular degrees of freedom of a C_α_-mapped model, designated as θ and α angles, which encompass α_1_ and α_2_ (Fig. 1a), thereby creating a pseudoramachandran plot that is suitable for a CG representation of a protein chain. In the CG topology, θ and α_2_ are, respectively, the angles between three subsequent C_α_-beads (*i_-1_*, i, and *i*_+1_) and the dihedral angle between four subsequent C_α_-beads (from *i_-1_* to *i*_+2_). ^43,44^

These angles reflect the values of φ and ψ in the all-atom space when converted into the C_α_-mapped framework and act as proxies of backbone conformational propensities for the CG model. They are particularly sensitive to long-range conformational backbone propensities as θ is an angle directly connected to two all-atom dihedrals associated with a C_α_ bead (ψ*_i_* and φ*_i_*), and the α angles reflect the contributions of four all-atom dihedral angles (e.g. ψ*_i_,* φ*_i_,* ψ*_i+1_* and φ*_i+1_* for α_2_).

As constraints naturally exist in all-atom protein backbones, not all values of α and θ are deemed possible.^43,44^ When projected into the α and θ space (Fig. 1c and S1), the NapshiftC_α_ dataset strongly populates α-helical or β-stranded regions (27.39 % coverage of the dataset) with a large proportion of residues surrounding the core of structured regions (72.49 % coverage) and generously allowed regions (98.75 % coverage). The variability of the training set is particularly important to sample both partially structured or transiently ordered segments, as well as rigorously folded α-helices and β-strands.

As our dataset spans the whole range of allowed CG angular space (Fig. 1c) NapshftC_α_ can learn to distinguish and generalise across the full spectrum of conformational variability, projected onto the α and θ angular spaces. Notably, the projection also shows a clear definition of disallowed regions: an important feature to minimise the sampling of near linear, unphysical conformers.

Once assessed the suitability of this dataset to describe the angular space of protein backbones, we then measured the ability of NapshiftC_α_ to predict backbone CS by monitoring the root mean squared error (RMSE) between experimental and computed CS. The results indicate that NapshiftC_α_ either matches or outperforms current all-atom CS predictors^45–48^ and yields similar results compared to the previously published all-atom Napshift.^49^ NapshiftC_α_ was also found to achieve a similar performance when compared to NapshiftCG: a previous ANN we tailored for the Martini3 CG model,^50^ despite the latter adopting two additional angles for its training, due to the presence of C_β_ beads in the CG mapping (Fig. 1d).

Confident of the level of accuracy of NapshiftC_α_ in modelling CS data, we then tested its ability to restrain MD simulations performed with the Calvados CG model^51^, by sampling the dynamics of a short peptide shown to fold into a α-helix in the presence of SDS micelles.^52^

Secondary shifts were calculated from the conformations sampled by Calvados in the absence and presence of NapshiftC_α_ restraints, and compared with the experimental NMR data, which show a strong signal reflective of a well-defined α-helix. Such a signal is well recovered by NapshiftC_α_-restrained Calvados, as it samples a strong α-helical ensemble as opposed to the unrestrained simulations (Fig. 1f) promoting a disordered ensemble and yielding random coil CS (Fig. 1e). A quantitative analysis of the cartesian and dihedral root mean square deviations (RMSD) confirms that NapshiftC_α_ minimises the average dihedral RMSD to ∼50° from ∼100° of the unrestrained simulations, and a mean C_α_ RMSD of ∼0.2 nm from the modelled α-helix (∼0.6 nm for the unrestrained ensemble). Notably, the C_α_ and dihedral RMSD distributions that NapshiftC_α_-restrained Calvados simulations yield, are much narrower than those obtained with unrestrained simulations, reflective of a well-defined and structured ensemble of conformations for the simulated peptide (Fig. 1f).

### Chemical shift restraints preserve experimental agreement with coarse-grained IDP dimensions

Given that NapshiftC_α_ restraints can recapitulate local structural properties of a simple C_α_-mapped peptide when applied to a force field not designed to reproduce secondary structure, we moved to understand how the application of CS would affect global properties of larger IDPs. We therefore performed Langevin dynamics simulations on a dataset of 65 proteins previously used to validate Calvados^51^ and part of the IDRome dataset.^53^ This set contains several members with a certain degree of predicted secondary structure, which in some cases is higher than 70%. For most proteins in this set, experimental NMR data are not available, requiring to generate synthetic CS using the PROSECCO server in combination with the prediction of residual secondary structure.^54^ The resulting dataset of proteins simulated with NapshiftC_α_ restraints yielded R_g_ values exhibiting a nearly full overlap with the values sampled by unrestrained Calvados simulations (ρ_c_=0.98, Fig. 2a-b). Unrestrained and NapshiftC_α_-guided Calvados ensembles reproduce equally well the experimental mean R_g_, reporting ρ_c_=0.93 and ρ_c_=0.92, respectively. In general, we found that the application of NapshiftC_α_ restraints introduces a systematic increase of R_g_ values relative to unrestrained simulations; this effect remains however minimal, even for systems exhibiting a high degree of secondary structure.

**Figure 2.**
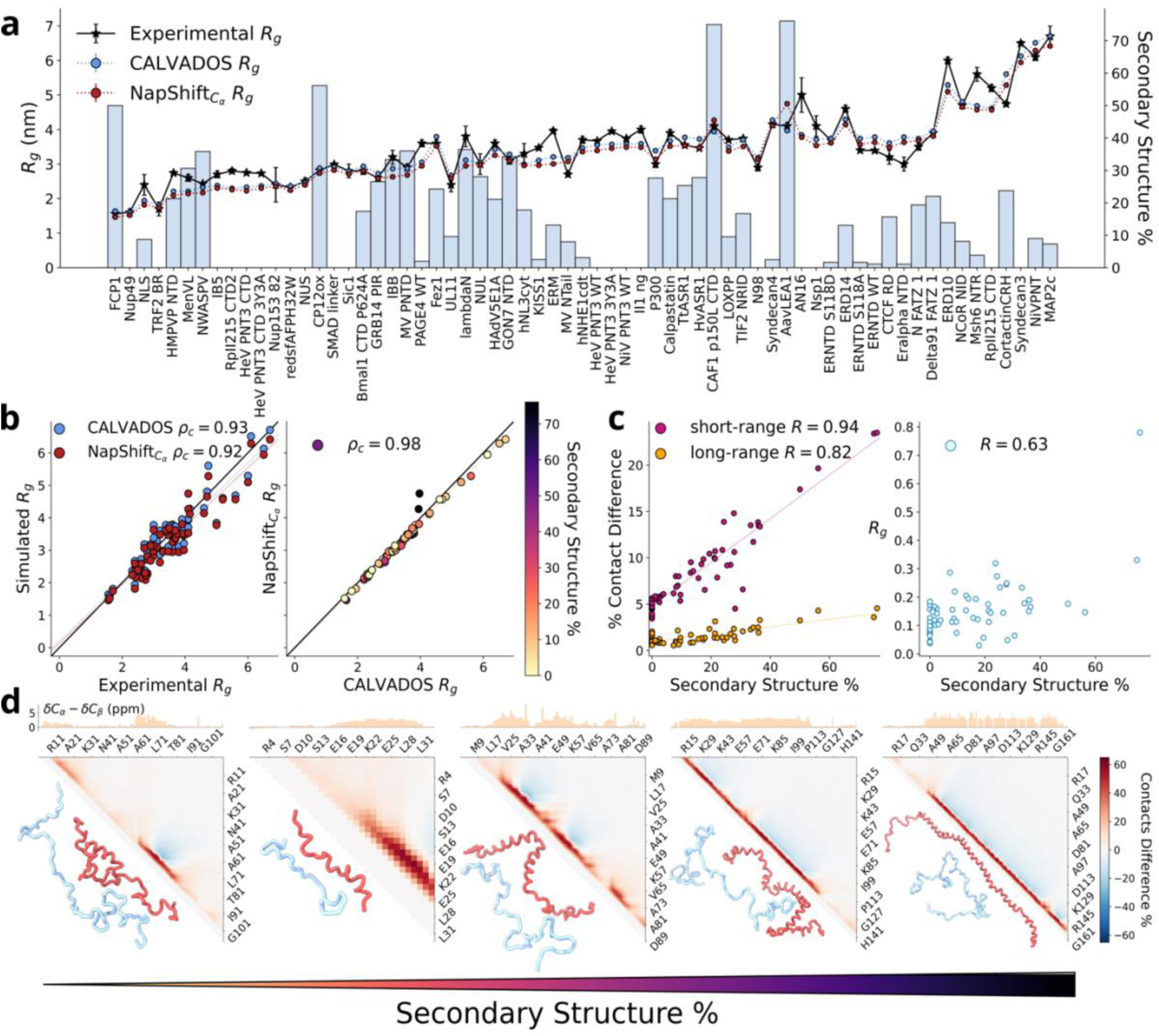
Validation of NapshiftC_α_ on single molecule IDPs. (a) Radius of gyration (R_g_) and secondary structure content of the validation dataset of Calvados within the IDRome dataset. The secondary structure percentage is represented by the blue bars, while the experimental and computed R_g_ are in black (experimental R_g_), and blue/red for Calvados/NapshiftC_α_. (b) Correlation between the experimental and computed R_g_ sampled by either Calvados (blue) or NapshiftC_α_ (red). (b) Correlation and concordance correlation coefficients (ρ_c_) between simulated and experimental radii of gyration (R_g_) (left) and between restrained and unrestrained simulations (right). (c) Correlation between the percentage of contacts difference between NapshiftC_α_ and Calvados and secondary structure (left) or R_g_ (right). (d) Contact maps of the five proteins in the dataset with the highest percentage of predicted secondary structure. From left to right: lambdaN, FCP1, CP12ox, C-terminal domain of CAF1p150L and AavLEA1 – see supplemental data excel spreadsheet for sequences. The contact maps show the difference in the number of intramolecular contacts sampled by NapshiftC_α_ compared to Calvados. Red and blue regions show increased and decreased contact percentage in restrained compared to unrestrained simulations. Representative conformations sampled in the simulations run with either Calvados (light blue) or NapshiftC_α_ (red), are shown in the lower triangle of each map.

We then investigated the link between intra-chain contacts and secondary structure content, to understand how the inclusion of secondary structure would influence bead-bead interactions.

We monitored the correlation of the observed contact difference seen between restrained and unrestrained simulations, and the predicted secondary structure percentage. The analysis indicated a strong correlation between secondary structure and short-range contacts with R=0.94 (Fig. 2c). This is expected, since secondary structure formation is associated with the establishment of local contacts leading to α-helical or β-strand moieties. This correlation, however, remains surprisingly strong (R=0.82) for contacts between beads separated by at least 15 bonds (Fig. 2c).

Given the potential relation between overall chain dimensions and secondary structure formation we also monitored the correlation between these two properties. In line with the minimal bias seen for R_g_ when NapshiftC_α_ restraints are applied, we did not find a significant correlation (R=0.63) between mean R_g_ and secondary structure (Fig. 2d). The analysis of the ensembles obtained from simulations of proteins with the highest secondary structure content shows that NapshiftC_α_-guided Calvados ensembles establish more short-range contacts focused along a contact map’s diagonal (Fig. 2e).

This finding suggests that the ensembles sampled by Calvados in the presence and absence of NapshiftC_α_ restraints are fundamentally divergent in contact space.

Besides the trends obtained from simulations carried out using synthetic CS, we also simulated proteins with experimental CS, where those were available.^55–60^ For those proteins, S4Pred^61^ is shown to over-estimate secondary structure (Fig. S2a), but the agreement with the experimental R_g_ remains unaffected (ρ_c_=0.92 for NapshiftC_α_-restrained Calvados simulations and ρ_c_=0.94 for unrestrained Calvados simulations), as the two simulation sets report a striking ρ_c_=0.99 (Fig. S2a-c). The application of CS restraints shows an improvement in the agreement between experimental and computed R_g_ (Fig. S2d), except for Calpastatin1, for which restraints slightly worsen the agreement.

It is noteworthy that the NMR data for Calpastatin1 (BMRB code: 15766) did not contain C_β_ shifts (Fig. S3 – y-axis for Calpastatin1), suggesting that the lack of experimental data could have likely contributed to a higher uncertainty in the determination of the ensemble. This confirms the importance to have exhaustive experimental data for an accurate ensemble estimation. In addition, as also seen for all the other proteins, the use of experimental CS generates ensembles that diverge in contact space: with unrestrained Calvados simulations which tend to sample longer range contacts than simulations restrained with NapshiftC_α_ (Fig. S3 – contact maps comparison).

Taken together, these data indicate that simulations carried out using either experimental or synthetic CS provide results of similar high quality, suggesting that the use of synthetic CS is a viable approach when experimental data is unavailable: as this is a common issue for IDPs.

Finally, to understand how CS restrained simulations would account for the behaviour of IDPs in condensates, we simulated the Calvados dataset of proteins with available experimental c_sat_ values.

We identify a systematic increase of c_sat_ values (Fig. S4), despite the two simulation sets retain a high degree of similarity (ρ_c_=0.96). It should be noted that for these proteins no secondary structure propensity was accounted for, as none was predicted by S4Pred, and it was not possible to assess the role of secondary structure on the qualitative or quantitative properties of the simulated condensates.

### NapshiftC_α_ rescues IDP conformational ensembles from overly linear conformations increasing smFRET agreement in secondary structure-containing regions

Given the observed differences in contact space between restrained and unrestrained simulations, we turned our attention to IDPs or IDRs for which more experimental information is available.

An increasing number of proteins have been recently investigated by both smFRET and NMR.^5,12,13,62^ We thus simulated the sex determining region Y box 2 transcription factor (Sox2), the high-affinity complex between prothymosin α and histone H1 (Protα-H1) and the full-length eukaryotic translation initiation factor 4b (eIF4b). For these systems, smFRET was performed on multiple labelled positions, providing a survey of intramolecular distances.^5,12,13,62^ These studies reported the comparison between computed and experimental mean FRET efficiencies (<E>), providing a direct mean of validation for computationally generated ensembles. More interestingly, while Sox2, Protα or H1 (aside from their well-structured domains) do not show any residual secondary structure, eIF4b is characterised by the formation of a α-helix between residues 361 and 407.^13^

The comparison of experimental and computed FRET efficiencies shows that unrestrained simulations performed with Calvados well-reproduce <E> across multiple labelled positions (Fig. 3a-c), and that a quantitative comparison between experimental and computed <E> yields good agreement, with ρ_c_=0.79, ρ_c_=0.83 and ρ_c_=0.80 for Sox2, Protα-H1 and eIF4b, respectively (Fig. 3a-c - correlation plots). The introduction of NapshiftC_α_ restraints, however, further improves this agreement to ρ_c_=0.86 and 0.87 for Sox2 and Protα-H1, respectively, with a reduced concordance correlation coefficient for eIF4b as ρ_c_ lowers from 0.8 to 0.76. This trend is however reversed when secondary structure is taken into consideration, as evidenced by an analysis of the four labelled positions indicated by triangles in Fig. 3c and conveying information on the protein region flanking and encompassing the α-helical segment. The comparison between experimental and computed <E> for the eIF4b regions 332-407, 332-457, 361-407, 361-457 strongly recovers the agreement, yielding a ρ_c_>0.8 (Fig. S5). Such a correlation is now comparable with that of unrestrained Calvados, but NapshiftC_α_ samples conformations that contain secondary structure, as guided by the experimental CS.

**Figure 3.**
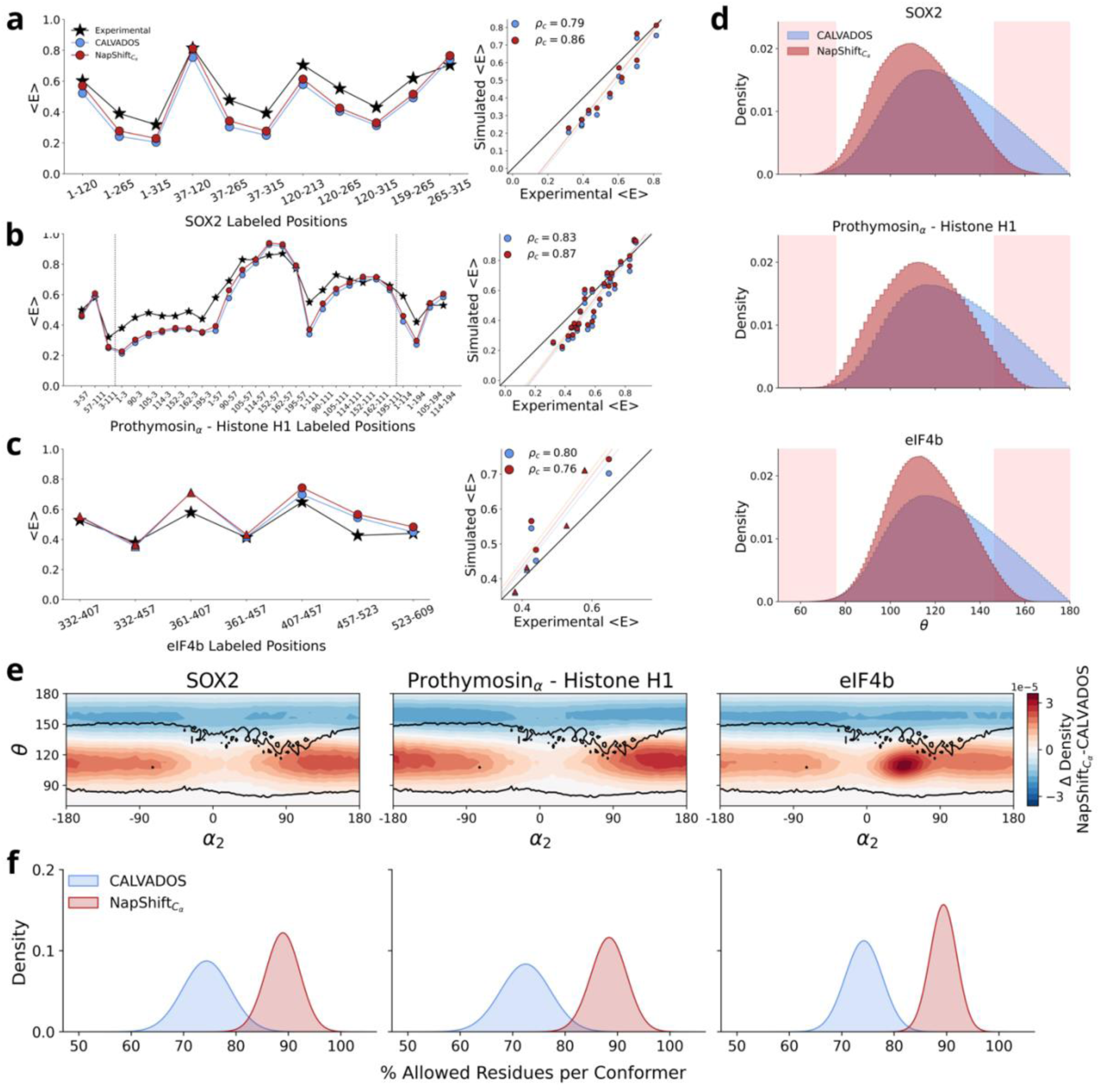
Agreement with smFRET spectroscopy for IDPs with different degree of secondary structure. (a-c) Mean FRET efficiencies (<E>) (blue – unrestrained simulations, red – restrained simulations, black experimental <E>) for labelled positions on Sox2 (a), the Protα-Histone H1 complex (b) and (c) eIF4B. In (c), the computed <E> for labelling positions that fall within the region of eIF4b flanking secondary structure are shown as triangles. Concordance correlation coefficients between experimental and computed <E> are shown on the right-hand side panels. (d) Density plots of the θ angles for the proteins in (a-c) simulated using either unrestrained (Calvados) or restrained (NapshiftC_α_). The red bands show disallowed regions of θ as described in the text. (e) Two-dimensional density plots of the θ angle as a function of dihedral α_2_ for the proteins in (a-c). The plots show the difference between restrained and unrestrained simulations, with red regions reporting on larger populations sampled by NapshiftC_α_ and blue regions on larger populations obtained from simulatiosn run with Calvados. Black lines show the boundaries of allowed angular pseudoramachandran regions as described in the text. (f) Density plots showing the percentage of residues falling within allowed pseudoramachandran regions in (e), from left to right: Sox2 (left), the Prothα-Histone H1 complex and eIf4b.

The same quantitative agreement of long-range experimental observables can thus reflect very differently shaped ensembles. In fact, when monitoring R_g_ and asphericity of these three proteins we observe no difference between restrained and unrestrained simulations (Fig. S6).

For the C-terminal region of eIF4b (residues 407 to 609) which shows the absence of secondary structure as indicated by low secondary CS, the agreement with experimental <E> is lower compared to Calvados, which has non-bonded interactions optimised to sample random coil conformations.

We then investigated how a similar agreement between restrained and unrestrained simulations would reflect in different conformational spaces. Relevantly, NapshiftC_α_ adopts a restricted bending potential (ReB), required when applying restraints to prevent the sampling of dihedral angles near 0° or 180°, as those are associated with inherent mathematical instabilities. This is in contrast with most CG force fields for IDPs which do not include angular terms in their Hamiltonian and might be naturally inclined to sample more linear conformations.^28^

We assessed the C_α_ backbone dihedral distributions sampled by unrestrained and NapshiftC_α_-restrained Calvados simulations, finding that the unrestrained ensemble is characterised by numerous conformations that explore disallowed θ values (Fig. 1c and S1), while NapshiftC_α_ restraints return a probability distribution of θ within allowed regions (Fig. 3d).

Having observed differences when monitoring θ, we asked if restrained and unrestrained simulations also sampled differences in the dihedral α space and plotted 2D distributions of θ vs. α_2_.

The analysis of the 2D plots in Fig. 3e shows that for all the proteins investigated, CS restraints drive the simulations to avoid the sampling of disallowed regions, while unrestrained simulations are prone to linearise conformers with θ consistently sampled well-above 150° over the whole α range (±180°). Moreover, the pseudoramachandran plot for eIF4B shows that NapshiftC_α_ improves the conformational space sampled by prompting a stronger α-helical sampling and suppressing overly linear conformers that unrestrained simulations tend to populate (Fig. 3e).

This consequently influences the whole ensemble, as we quantify the percentage of residues, per sampled conformer, that fall in disallowed regions of the pseudoramachandran space shown in Fig. 3e. This analysis returns non-overlapping distributions for restrained and unrestrained simulations, with unrestrained simulations sampling conformers with only approximately 70% of residues (distribution median), compared to 90% for restrained simulations.

### Using CS restraints suggests an active role of structured elements in defining condensate properties

Given that eIF4b contains a α-helix (Fig. 4a) and can organise in condensates driven by its aromatics-containing DRYG domain, we investigated how the presence of CS restraints and particularly, the sampling of a α-helical region, would affect the ability of the protein to organise in a condensate. In a previous work, we used unrestrained Calvados to simulate eIF4b in condensates, obtaining a good agreement with experimental mean FRET efficiencies.^13^ Here we monitored the ability of unrestrained and restrained simulations of eIF4b condensates to reproduce experimental c_sat_ faithfully, as secondary chemical shifts (Fig. 4b) had been experimentally obtained by Swein et al.^13^

**Figure 4.**
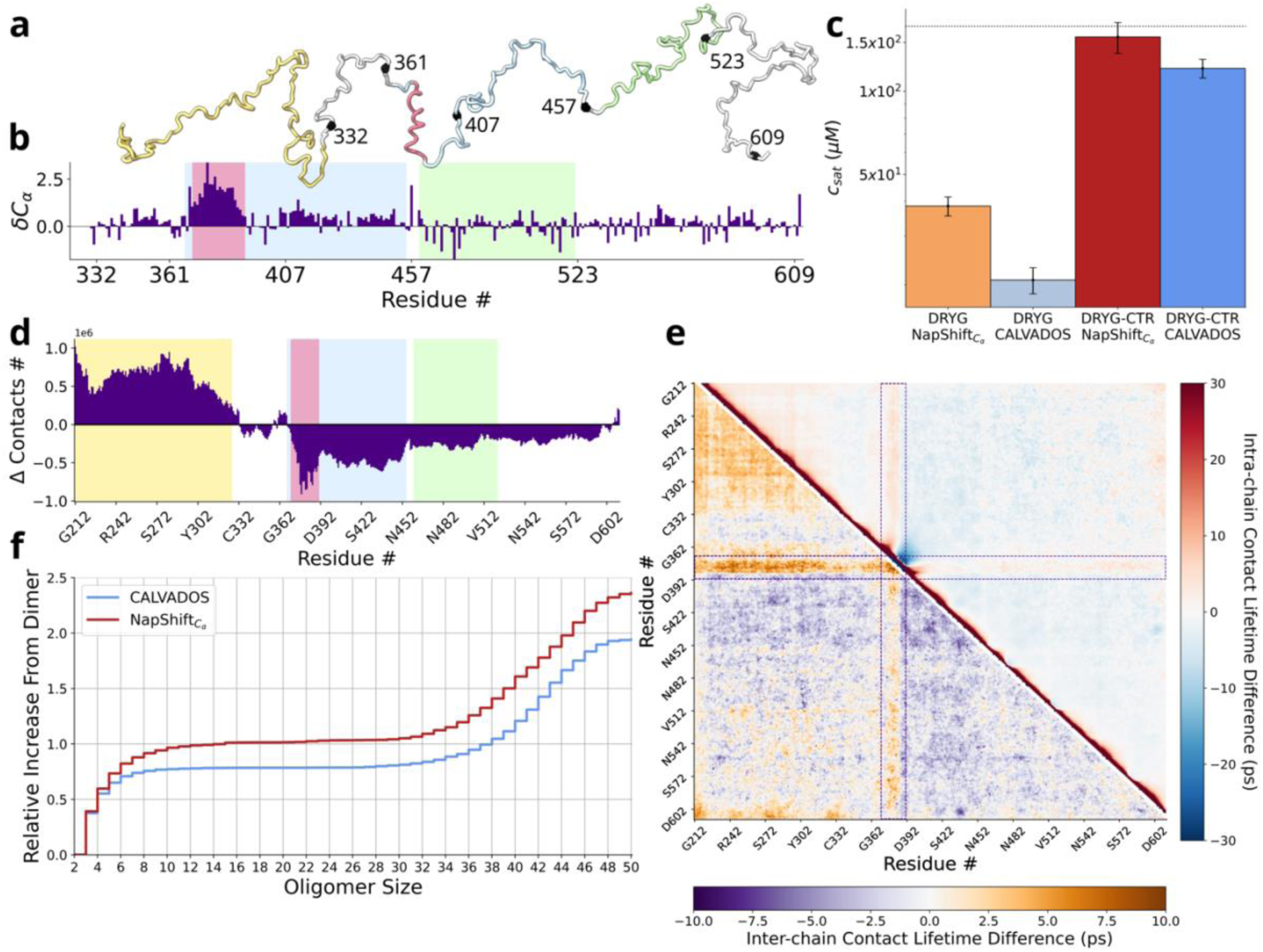
Effects of secondary structure from restrained simulations of eIF4b in condensates. (a) Representative conformation of eIF4B highlighting the protein’s subdomains. The DRYG-rich domain (residues 213-332), the helical region (residues 361-407), the RE-rich domain (residues 367-455) and the proline-rich (residues 460-522) domains are coloured yellow, red, cyan and green, respectively. Interdomain segments and C-terminal end of the protein are coloured in grey. The black spheres correspond to the positions of the experimental FRET labels. (b) Experimental secondary chemical shifts for eIF4b reported by Swain et al.^13^ (c) Computed c_sat_ for the DRYG domain alone or for the entire eIF4b (DRYG-CTR), sampled via restrained (yellow and red) or unrestrained (light and dark blue) simulations. (d) Differences between the number of contacts made in the condensate as a function of eIF4b sequence. The difference was calculated by subtracting the number of contacts sampled by NapshiftC_α_ from those sampled by Calvados. Positive and negative values show more and less contacts made in NapshiftC_α_ simulations. (e) Top triangle - intra-chain contact differences between simulations performed with NapshiftC_α_ and Calvados. Red and blue regions show NapshiftC_α_ making more and less contacts compared to Calvados, respectively. Bottom triangle – inter-chain contact lifetime differences between simulations performed with NapshiftC_α_ and Calvados. Orange and purple regions show NapshiftC_α_ making longer-lived or shorter-lived contacts compared to Calvados, respectively. (e) Cumulative distribution functions of the oligomeric species found in the condensate simulated via restrained (red) or unrestrained (blue) simulations. The fraction of the different oligomers are shown with respect to the smaller oligomer (2-mer).

For both simulation sets we observed significantly reduced c_sat_ values when only the DRYG domain is considered (Fig. 4c), with ensembles that are visually similar (Fig. S7). The considerably lower c_sat_ for the DRYG-only system is unsurprising, as experiments show that the DRYG domain is an active driver of the phase separation of eIF4b.^13^ We however observe differences in c_sat_ (beyond experimental uncertainty) between restrained and unrestrained simulations (Fig. 4c) when comparing the computed and experimental c_sat_ of the whole protein (DRYG-CTR). In this case, NapshiftC_α_-restrained Calvados simulations were found to better match experimental c_sat_ (Fig. 4c – dashed line) obtained upon fitting experimentally measured c_sat_ as a function of the ionic strength through a binding isotherm (Fig. S8).

The differences observed between restrained and unrestrained simulations could be interpreted in two ways: either as a direct consequence of the CS restraints regardless of the α-helix sampling (random coil CS), or as arising from an active role of the α-helix within the simulated condensate.

To address these two interpretations, we monitored the intermolecular contact difference as a function of the eIF4b sequence and found that (i) NapshiftC_α_ simulations sample a higher number of contacts occurring within the DRYG domain and (ii) the α-helical region in eIF4b shows the strongest difference in contact space when compared to the unrestrained simulations (Fig. 4d). To quantitatively monitor interactions within the condensate, we also calculated intra- and inter-chain contact lifetimes, to understand if longer contact lifetimes would have led to the observed difference in c_sat_. While most intra-chain contact lifetime differences are expectedly concentrated along the map’s diagonal, NapshiftC_α_-restrained Calvados simulations also show considerably off-diagonal long-range inter-chain contacts between DRYG domains and especially along the α-helical region, which is found to be a hotspot for contact formation (Fig. 4e). These conformational properties affect c_sat_ and the previously observed ability of eIF4b to condense via the formation of smaller oligomers.^13^ In fact, simulations with NapshiftC_α_ restraints report a higher population of smaller oligomers compared to unrestrained sampling (Fig. 4f). Taken together these analyses therefore indicate an active role of the helical moiety in modulating the dynamics of the DRYG and C-terminal domains flanking it, as the observed effects would ultimately have an impact on c_sat_ .

## Discussion

We here present the training and application of an ANN that we call NapshfitC_α_, and find that it can effectively predict CS in over-simplified (C_α_-only) CG models. We show the applicability of NapshfitC_α_ as a restraining tool for MD simulations, to sample IDP dynamics in both single chains and condensates. Nuclear Overhauser Effect (NOE) distance restraints, which inform on the packing of structured domains,^63^ were also employed when sampling the dynamics of folded protein domains alongside intrinsically disordered regions (IDRs). Such a combination of CS and NOEs was found to convey both local and global bead re-arrangements allowing to effectively simulate proteins containing both order and disorder. Moreover, in the absence of experimental data, CS predictive tools such as PROSECCO^64^ in combination with accurate secondary structure predictions, were shown to be valuable sources of synthetic NMR data to be employed in accurate restrained simulations, as tested on a large dataset of proteins.

Numerous attempts to computationally describe IDP behaviour have focused on the development of several C_α_-only potentials strongly focusing on the optimisation of non-bonded interactions through different theoretical paradigms.^18,20,22,23,25,28,51,65–67^ The parametric space of these models, being largely confined to modified Lennard-Jones potentials, tends to neglect local environmental effects and therefore, by design, does not explicitly account for the formation of secondary structure, which is recognised as a dominant functional element of IDPs.^68^ The unstructured nature of IDPs stresses the importance of sequence-to-function relationships in cellular contexts. ^69^ This has triggered the notion that the function of IDPs can be mostly described on an ensemble-to-function basis, ^70^ reflected through the ability of simulations to correctly sample global conformational features of IDPs proxied by R_g_ or longer-range distances within a chain, as obtained from SAXS and smFRET spectroscopy.^6,8,9^

We show that it is possible to effectively reproduce secondary structure propensity in coarse graining through the direct introduction of NMR observables: even when coarse graining is as minimalist as one bead-per-residue. The integration of CS restraints into these CG simulations was found to have a strong effect on the ensembles in local and global contact space, although it did not perturb the overall agreement with long-range conformational properties dictated by experimental R_g_ and <E>. These properties are indeed well-described by the available non-bonded potentials^18,71^: carefully optimised to match global characteristics of IDPs. Global properties are however poorly informative of local environments, as evidenced in the diverging ensembles yielded by restrained and unrestrained simulations in spite of the observed agreement in R_g_ and c_sat_.

The introduction of NMR observables directly informing backbone dihedrals can ameliorate the sampling of residues falling in disallowed regions of the CG modelling space, through the statistical learning of the ANN and the application of restraints that have learnt the boundaries between allowed and disallowed backbone angles and dihedrals. In the absence of angular and dihedral potentials, we find that Calvados is prone to sample linear-like conformations, which could be also the case for other CG models lacking angle or dihedral potentials. NapshiftC_α_ can avoid this while improving ensemble accuracy by matching both R_g_ and mean FRET efficiencies from SAXS or smFRET, as well as sampling secondary structure as indicated by the collected NMR data. This is particularly relevant if data from CG simulations of IDPs are then used to train ensemble predictors aiming at generating accurate IDP conformations^72^, since those would reflect the capacity of a force field to correctly sample single conformers beyond global chain properties. Restrained simulations thus improve our understanding of ensemble-function relationships in IDPs, useful to study IDPs and IDRs in cellular environments or to design intrinsic disorder with desired conformational properties, as previously suggested.^53,73^ For these tasks, an accurate interpretation of ensembles and a quantitative match of observables is crucial.

Indeed, the comparison of restrained and unrestrained simulations also informs how secondary structure can influence not only local properties within short-range distances, but at some extent also longer-range contacts, concentration of saturation and metastability profiles in condensates, as in the case of eIF4b: a translation factor featuring a α-helical region and shown to phase separate through the formation of metastable oligomers.^13^. Relevantly, we find that in condensates the helical segments of eIF4b can play a significant role in defining the concentration of saturation (c_sat_) and that secondary structure has an influence on the distribution of large-size oligomers shaping the phase-separation landscape of the protein.^13^ This finding indicates that for functional traits more heavily mediated by short-range contacts, such as short-linear motifs (SLiMs) interacting with structured targets, sampling local secondary structure is paramount, and NapshiftC_α_ is expected to provide a significant improvement in studies using either synthetic or experimental NMR data. The adoption of synthetic CS is however strictly dependent on the quality of the predictor generating CS data, as we show that, while in many cases predictors suggest CS are in qualitative agreements with experiments, they can over or under-estimate secondary shifts.

Being trained on a large dataset, NapshiftC_α_ shows exquisite sensitivity in its ability to tie NMR observables to computed conformational space and to obtain reliable ensembles. Thus, the capacity to access NMR data repositories is crucial for the successful development of methods able to investigate the conformational properties of IDRs, as it has been suggested for structured proteins.^74^ Notably, the reliability of experimental NMR data is recently being investigated in relation to the validity of conformational ensembles.^75^ Overall, with access to robust experimental NMR data, NapshiftC_α_ can be tailored to any C_α_-mapped CG force field for IDPs, leading to a more experimentally-informed and physically grounded exploration of IDP behaviour; achieved by grasping how secondary structure in IDPs can be functionally important in both single chain and condensates.

## Acknowledgements

M.C. acknowledges the Riddet Institute for supporting part of this work through a PhD scholarship. D.M. acknowledges the support of the Riddet Institute and the Maurice Wilkins Centre partially funding the research activities presented in this work.

## Author Contributions

M.C. trained the ANN, implemented the framework within OpenMM, ran simulations and analysed data; A.D.S. and D.M. conceived and designed the study; D.M. and K.T. jointly supervised M.C.; C.B., V.M. and A.D.S. advised M.C. on several aspects of the work; D.M. wrote the manuscript with the contribution of all the authors.

## Data and code availability

The excel spreadsheet named “SupplementalData.xlsx” provides the following data: the spreadsheet named “Dataset” provides the list of PDB files for the train, validation and test sets. The spreadsheet named “Demo System” contains simulation conditions, peptide sequence and DOIs for the peptide presented in Fig. 1. The spreadsheet named “Calvados validation” provides a table with information of the 65 protein systems presented in Fig. 2. The spreadsheet named “Calvados slab” provides a table with information of the 32 protein systems presented in Fig. S3.

The spreadsheet named “FRET systems” provides a table with simulation conditions, sequence and DOIs of the eIF4b, Sox2, prothymosinα and Histone H1 complex presented in Fig. 3 and 4. The spreadsheet named “eIF4B Condensate” provides a table with simulation conditions, sequence and DOIs of the eIF4b system, simulated in the condensed phase and presented in Fig. 5.

The spreadsheet named “ADF statistics” provides Augmented Dickey-Fuller (ADF) coefficients calculated to assess stationarity of all the simulations pertaining to Fig. 1 and 3.

Napshift plugins for OpenMM will be made available upon publication of the peer-reviewed article.

## Methods

### Data preparation, training and restraints of Napshift***C_α_***

A set of 3,236 NMR structures of proteins was gathered from the Protein Data Bank (PDB), with associated CS sourced from the BioMagnetic Resonance Data Bank (BMRB).^76^ When multiple models were provided within a single PDB file, only the first (that of the lowest energy) was considered.

Of these structures, 2,986 were used for training and validation of the CS prediction model, while 250 were reserved for testing. CS values more than 3 standard deviations from the average value reported for their residue-atom type in the BMRB were considered inaccurate and discarded.

During pre-processing, raw CS values were converted to secondary CS by subtracting their random-coil CS, as calculated by CamCoil.^77^

An Artificial Neural Network (ANN) was trained to predict secondary CS for the 6 backbone atoms (N, C, C_α_, C_β_, H_α_ H) from sequence and structure information of a protein represented by C_α_ atoms alone. Input vectors to the model combined information about local geometry and amino-acid type.

For each protein residue i, we describe calculate θ as follows:

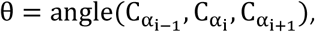

with α_1_ and α_2_ being the following torsions:

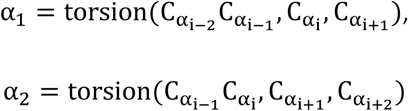

We represent these angles using their (sin θ, cos θ) pair. An embedding of amino-acid type was extracted from the BLOSUM62 matrix. In combination with the standard 20, two additional amino acid types were used to represent oxidized cysteine (CYO) and cis proline (PRC). This yields a vector of 28 values for each residue: 22 for amino acid type, and 3x2 for angular geometry. In practice, we concatenate a residue’s input vector with those of the residues before and after in sequence to produce a tri-peptide representation ending up being a vector of 28x3 values. The ANN was composed of an input layer (size 84), a hidden layer (size 26) equipped with the Exponential Linear Unit (ELU) activation function^78^, and an output layer (size 6) with a linear activation function appropriate for regression.

The size 6 output layer predicts the secondary CS for the 6 backbone atoms simultaneously. The ANN was trained on a set of 272,984 tripeptides extracted from 2,687 training structures.

Mean Squared Error (MSE) was used as the loss function and the ADAM optimiser with a learning rate of 0.001 was used to adjust model parameters. An early-stopping criterion assessed after each epoch on 32,076 tripeptides from 299 held-out validation structures helped to prevent overfitting by halting training if the validation loss failed to improve after 5 epochs. The best model obtained after early-stopping was tested on 28,107 tripeptides extracted from the 250 test structures.

The discrepancy between simulated CS predicted by the ANN and those measured by NMR experiments yields the following harmonic restraint potential:

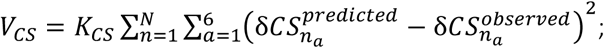

where 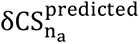 is the secondary CS predicted by the ANN for atom a of residue n, 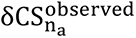 is the experimentally observed value, and K_CS_ is the force constant of the restraint potential.

In practice, the CS restraint potential is ramped up by linearly increasing K_CS_ from 0 to 25 over 25,000 steps. The CS restraint potential has been implemented as a plugin for the OpenMM simulation engine version 8, with GPU support.^79^ Numerical instabilities from the angular potentials were resolved by introducing a restricted bending potential (ReB), like that proposed in Bulacu et al.^80^, as follows:

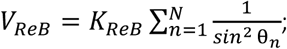

where K_ReB_ is the force-constant of the ReB potential.

As with the CS restraints, we ramp up this potential by linearly increasing K from 0 to 1 over 25,000 steps.

To recover protein tertiary packing, we employ distance restraints derived from NOESY experiments. Given atoms a_i_, a_j_ with an NOE-derived interatomic distance d_ij_, and which map to C_α_ beads A_i_, A_j_ respectively, their coarse-grained NOE distance D(A_i_, A_j_) is calculated as:

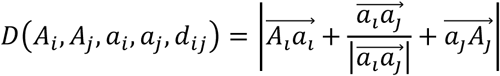

When multiple atoms contribute to the same NOE signal, we condense their inter-atomic distances to a single coarse-grain distance, and calculate:

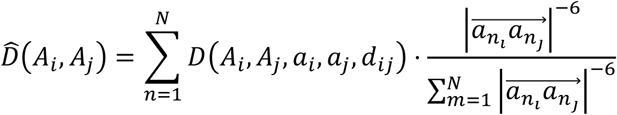

After condensing signals from multi-pair atomistic NOEs into a single coarse-grain distance, there may still be multiple experimental signals which map onto the same C_α_ bead pair A_i_, A_j_. Here, the resultant coarse-grain NOE distance is calculated as the average coarse-grain NOE distance for that C_α_ bead pair:

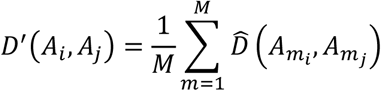

Having translated atomistic NOE data into coarse-grain distances, we employ a distance restraint potential similar to that described in Torda and van Gusteren^81^:

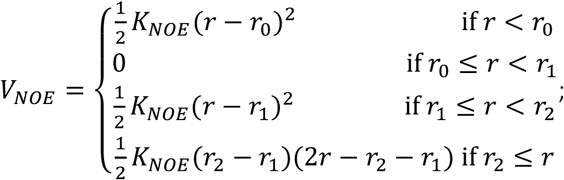

where r is the distance between two C_α_ beads and K_NOE_ is the force constant of the NOE restraint potential. This potential is composed of a harmonic regime, a flat-bottom regime between r_0_and r_1_ to account for uncertainty in the measurement, and a linear regime beyond r_2_.

Where r_0_ and r_2_ are missing from experimental data, we set r_0_ = 0 and r_2_ = r_1_ + 0.5 nm. Restraining forces are applied in a time-averaged fashion^82^ as follows:

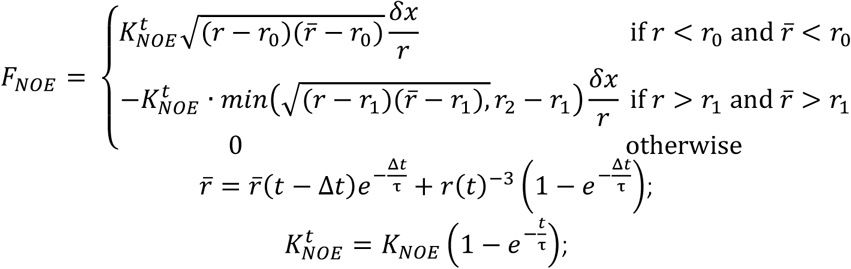

where δx is the direction vector between the two C_α_ beads, t is the current simulation step, and τ = 0.05 ps is the decay time for the exponential running average. As with the CS restraint potential, this NOE restraint potential has been implemented as a plugin for OpenMM.

The complete set of restraint potentials (CS, restricted bending - ReB, and NOE) are added on top of the underlying forcefield to produce a total potential energy given by:

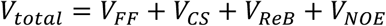

Prediction of protein secondary-structure propensity from sequence was performed by S4Pred, which produced for each residue a prediction of the probability of each secondary structure type (α-helix, β-sheet and coil). For proteins with NMR data available, we used SSP^83^, which calculates similar probabilities from secondary CS.

To produce synthetic CS sensitive to the variable structure/disorder propensity in our dataset, we used the PROSSECO^64^ server to predict fully disordered CS (from sequence alone) and structured CS (from sequence and S4Pred predictions). For each residue, we combine these two sets of CS by weighting them according to the residue’s S4Pred-predicted structural propensity:

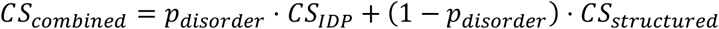

### Langevin simulations

Coarse-grained Langevin dynamics simulations were carried by OpenMM 8.2.0^79^, using the Langevin integrator with a time step of 10 fs and friction coefficient of 0.01 ps^-1^, under periodic boundary conditions. Simulations were performed using the CALVADOS2 force field.^25^ Each residue was modelled as a single bead, mapped on the C_α_ atom. A harmonic potential maintains bonds between beads:

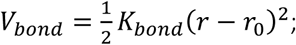

where the force constant k=8033 kJ mol^−1^ nm^−2^ and the equilibrium bond distance r_0_=0.38nm. Non-bonded potentials were given by a truncated-shifted Ashbaugh-Hatch potential:

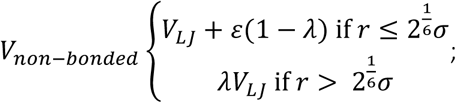

where ɛ = 0.8368 kJ mol^−1^, r_c_ = 2 nm, and U_(LJ)_ is the Lennard-Jones potential. The σ and λ parameters were set as in CALVADOS2.^25^

A Debye-Huckle potential was used to define salt-screened electrostatic interactions:

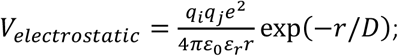

where q is the charge of an amino acid. We specifically set q=-1 for aspartate and glutammate, q=+1 for lysine and arginine, and q=+0.5 for protonated histidine at pH 6, or q=0 otherwise), *e* is the elementary charge. The Debye length 𝐷 for a given salt concentration 𝑐_𝑠_ is calculated as:

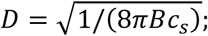

where 𝐵(𝜀_𝑟_) is the Bjerrum length.

The temperature-dependent dielectric constant 𝜀_𝑟_ is given by:

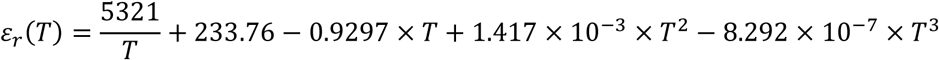

Electrostatic interactions were treated through a shifted-cutoff potential, where the cutoff r_c_ was set at 4 nm.

We simulated 65 IDP sequences not included in the CALVADOS training set were simulated at 310 K at an apparent ionic strength of 150 mM.

The initial coordinates were modelled in Archimedes’ spiral with a spacing of 0.38 nm C_α_ beads and placed in a cubic box of length 0.38 x (N-1) + 4 nm, where N is the number of protein residues. After 25000 steps, during which the CS restraints and the ReB potential were gradually ramped up, coordinates were reported every Δt = 3N^2^ fs if N> 150 and Δt = 70 ps otherwise.

SOX2, eIF4B, and Protα-H1 were simulated for 10 μs, recording frames every 1000 steps.

These were simulated with temperatures and salt concentrations reflecting their respective experimental conditions; 293 K at 192 mM salt, 298 K at 165 mM salt, and 300 K at 165 mM salt.

For these systems, we use experimental CS for each residue where these are available, and random coil CS predicted by CamCoil^77^ otherwise. Specifically, CS for the C-terminal region of eIF4B, disordered regions of SOX2, and full length Protα were sourced from BMRB entries 51957, 51964, and 27216, respectively. For the systems with structured domains - SOX2 and Protα-H1 - we employed distance restraints from NOE data. Initial structures, NOE distances, and CS values for the globular domains of SOX2 and Histone H1 were taken from PDB IDs 2LE4 and 6HQ1, respectively.

Direct coexistence simulations were performed in a cuboidal box of dimensions 17 x 17 x 300 nm for Ddx4_WT_, 25 x 25 x 200 nm for eIF4B, and 15 x 15 x 150 nm for all other systems. 100 chains were simulated for each system except eIF4b, for which we simulated 50 chains.

Simulations of the CALVADOS validation set used temperatures and salt concentrations derived from each system’s experimental conditions, while eIF4B was simulated at 293K at 117mM salt concentration.

The starting conformation for each chain was generated as an Archimedes’ spiral.

These spirals were arranged along the z-axis with a spacing of 1.47 nm. To produce starting conformations for production, we follow the method described in Dima and Thirumalai^84^, whereby equilibrium runs of 2M (million) steps were performed with an external force pulling each chain towards the box center, forming a condensate. Systems were then simulated for 5 μs, reporting every 250000 steps (or for 10 μs reporting every 10,000 steps in the case of eIF4b). For all analysis, the first 1 μs was discarded as equilibration time. All simulations were performed in replicates of three.

### Simulation analysis

For each frame, the radius of gyration, 𝑅_𝑔_, was calculated as:

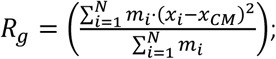

where 𝑁 is the sequence length, 𝑚 is the mass of bead 𝑖, 𝑥_𝑖_ is the position of bead 𝑖, and 𝑥_𝐶𝑀_ is the position of the protein’s center of mass. The asphericity δ is calculated as:

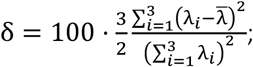

where λ_𝑖_is the 𝑖^𝑡ℎ^ eigenvector of the inertia tensor, and λ̅ is the mean of λ_1_, λ_2_, λ_3_.

Agreement between 𝑅_𝑔_ measured in experiments and calculated from simulations was quantified by the concordance correlation coefficient ρ_𝑐_:

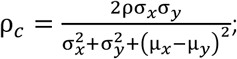

where μ and σ are the means and standard deviations of the datasets x and 𝑦, and ρ is the pearson correlation coefficient of x and 𝑦.

### Contacts analysis

For single chain simulations, C_α_ beads were considered in-contact when the distance between them was less than 2 nm. Contact matrices were calculated as the fraction of frames for which a given residue-residue pair was formed. Contact difference matrices were calculated by subtracting one contact matrix from another. A single metric for comparison between ensembles in contact-space was obtained by taking the RMSD between two contact matrices. This was calculated considering only residue-residue pairs within and beyond 15 amino acids of one-another to produce measurements of short-range contact-difference and long-range contact-difference respectively.

### Concentration of saturation

For calculations of c_sat_, the slab was centred in the box, and a density profile was calculated and used to estimate the dense and dilute phases, according to the method described in Tesei et al.^51^ An estimate for the experimental c_sat_ of eIF4b at 117mM was obtained by fitting a dose-response curve to the data reported in Swain et al.^13^

### FRET

smFRET transfer efficiencies were calculated from simulated distances via the Förster equation:

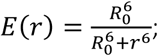

where 𝑟 is the distance between two labelled residues, and the Förster radius 𝑅_0_ is set according to the dye-pair used in the smFRET experiments: 5.9 nm, 6.0 nm, and 5.4 nm for eIF4b, SOX2, and Protα-H1, respectively. While FRET dyes were not explicitly represented in simulations, the effect of these dyes on the resulting ensembles has been shown to be negligible.^5,12,62^

### Angles and dihedrals

Distributions of α and θ were gathered from the NapShiftC_α_ training dataset and used to create a 2D histogram with bins with a width of 3°. From this, the ‘generously allowed’ region was defined as all bins populated by at least 10 α/θ pairs, as in Kleywegt.^43^ Similar distributions of α and θ were produced from simulations of eIF4B, SOX2, and Protα-H1. For a given conformation, the fraction of allowed residues was calculated as the number of residues whose α/θ pair fell within the ‘generously allowed’ region calculated from the NapShift dataset, divided by the sequence length.

### Contact Lifetimes

Contact lifetime analysis of eIF4B direct-coexistence simulations employed the transition-based definition described in Galvanetto et al.^85^, whereby a contact is considered formed when the C_α_-C_α_ distance drops below r_0_ and considered broken at the next time that this distance rises above r_1_.

We set r_0_ = 1.0 nm, and r_1_= 2.0 nm. Average lifetimes of each residue–residue contact were calculated by dividing the total bound time by the total number of breaking events for that contact. These were condensed to a single lifetime for each interchain and intrachain residue-residue pair by taking the average contact lifetime for that inter/intra residue pair across all eIF4B chains (ignoring pairs which did not form this contact). Interchain contact counts were calculated by summing the number of occurrences of each interchain residue-residue contact over all chains.

### Cluster analysis

For each frame, as in the gmx clustsize algorithm, chains were assigned to the same cluster on the condition that they share at least one pair of beads at most 1 nm apart. A cumulative distribution of cluster sizes was obtained and normalized to the number of dimer (2-mer) clusters. Simulated conformations were visualized using VMD version 1.9.4^86^ and ChimeraX version 1.8.^87^

## References

1. Holehouse, A. S. & Kragelund, B. B. The molecular basis for cellular function of intrinsically disordered protein regions. Nat Rev Mol Cell Biol 25, 187–211 (2024).

2. Babu, M. M., Kriwacki, R. W. & Pappu, R. V. Versatility from protein disorder. Science (1979) 337, 1460–1461 (2012).

3. Aznauryan, M. et al. Comprehensive structural and dynamical view of an unfolded protein from the combination of single-molecule FRET, NMR, and SAXS. Proc Natl Acad Sci U S A 113, E5389–E5398 (2016).

4. Schneider, R. et al. Visualizing the molecular recognition trajectory of an intrinsically disordered protein using multinuclear relaxation dispersion NMR. J Am Chem Soc 137, 1220–1229 (2015).

5. Borgia, A. et al. Extreme disorder in an ultrahigh-affinity protein complex. Nature 555, 61–66 (2018).

6. König, I. et al. Single-molecule spectroscopy of protein conformational dynamics in live eukaryotic cells. Nat Methods 12, 773–779 (2015).

7. Schuler, B., Soranno, A., Hofmann, H. & Nettels, D. Single-Molecule FRET Spectroscopy and the Polymer Physics of Unfolded and Intrinsically Disordered Proteins. Annu Rev Biophys 45, 207–231 (2016).

8. Metskas, L. A. & Rhoades, E. Single-molecule FRET of intrinsically disordered proteins. Annu Rev Phys Chem 71, 391–414 (2020).

9. Naudi-Fabra, S., Tengo, M., Jensen, M. R., Blackledge, M. & Milles, S. Quantitative Description of Intrinsically Disordered Proteins Using Single-Molecule FRET, NMR, and SAXS. J Am Chem Soc 143, 20109–20121 (2021).

10. Liu, Z. H., Tsanai, M., Zhang, O., Forman-Kay, J. & Head-Gordon, T. Computational Methods to Investigate Intrinsically Disordered Proteins and their Complexes. http://arxiv.org/abs/2409.02240 (2024).

11. Mercadante, D. et al. Kirkwood-Buff Approach Rescues Overcollapse of a Disordered Protein in Canonical Protein Force Fields. Journal of Physical Chemistry B 119, (2015).

12. Heidarsson, P. O. et al. Release of linker histone from the nucleosome driven by polyelectrolyte competition with a disordered protein. Nat Chem 14, (2022).

13. Swain, B. C. et al. Disordered regions of human eIF4B orchestrate a dynamic self-association landscape. Nat Commun 15, 8766 (2024).

14. Milles, S. et al. Plasticity of an Ultrafast Interaction between Nucleoporins and Nuclear Transport Receptors. Cell 163, (2015).

15. Childers, M. C. & Daggett, V. Validating Molecular Dynamics Simulations against Experimental Observables in Light of Underlying Conformational Ensembles. Journal of Physical Chemistry B 122, 6673–6689 (2018).

16. Brotzakis, Z. F., Zhang, S., Murtada, M. H. & Vendruscolo, M. AlphaFold prediction of structural ensembles of disordered proteins. Nature Communications 2025 *16:1* 16, 1–9 (2025).

17. Dinic, J., Marciel, A. B. & Tirrell, M. V. Polyampholyte physics: Liquid–liquid phase separation and biological condensates. Curr Opin Colloid Interface Sci 54, 101457 (2021).

18. Dignon, G. L., Zheng, W., Kim, Y. C., Best, R. B. & Mittal, J. Sequence determinants of protein phase behavior from a coarse-grained model. PLoS Comput Biol 14, (2018).

19. Dignon, G. L., Zheng, W., Best, R. B., Kim, Y. C. & Mittal, J. Relation between single-molecule properties and phase behavior of intrinsically disordered proteins. Proc Natl Acad Sci U S A 115, 9929–9934 (2018).

20. R. Tejedor, A. et al. Chemically Informed Coarse-Graining of Electrostatic Forces in Charge-Rich Biomolecular Condensates. ACS Cent Sci 11, 302–321 (2025).

21. Zerze, G. H. Optimizing the Martini 3 Force Field Reveals the Effects of the Intricate Balance between Protein-Water Interaction Strength and Salt Concentration on Biomolecular Condensate Formation. J Chem Theory Comput 20, 1646–1655 (2024).

22. Joseph, J. A. et al. Physics-driven coarse-grained model for biomolecular phase separation with near-quantitative accuracy. Nat Comput Sci 1, 732–743 (2021).

23. Wang, L., Brasnett, C., Borges-Araújo, L., Souza, P. C. T. & Marrink, S. J. Martini3-IDP: improved Martini 3 force field for disordered proteins. Nature Communications 2025 16:1 16, 1–14 (2025).

24. von Bülow, S. et al. Prediction of phase-separation propensities of disordered proteins from sequence. Proceedings of the National Academy of Sciences 122, e2417920122 (2025).

25. Tesei, G. & Lindorff-Larsen, K. Improved predictions of phase behaviour of intrinsically disordered proteins by tuning the interaction range. Open Research Europe 2, 94 (2023).

26. Thomasen, F. E., Pesce, F., Roesgaard, M. A., Tesei, G. & Lindorff-Larsen, K. Improving Martini 3 for disordered and multidomain proteins. J. Chem. Theory Comput. 18, 2033–2041 (2022).

27. Rizuan, A., Jovic, N., Phan, T. M., Kim, Y. C. & Mittal, J. Developing Bonded Potentials for a Coarse-Grained Model of Intrinsically Disordered Proteins. J Chem Inf Model 62, 4474–4485 (2022).

28. Hu, Z., Sun, T., Chen, W., Nordenskiöld, L. & Lu, L. Refined Bonded Terms in Coarse-Grained Models for Intrinsically Disordered Proteins Improve Backbone Conformations. Journal of Physical Chemistry B 128, 6492–6508 (2024).

29. Camacho-Zarco, A. R. et al. NMR Provides Unique Insight into the Functional Dynamics and Interactions of Intrinsically Disordered Proteins. Chem Rev 122, 9331–9356 (2022).

30. Schiavina, M. et al. Intrinsically disordered proteins studied by NMR spectroscopy. J Magn Reson Open 18, 100143 (2024).

31. Liu, Z. H., Tsanai, M., Zhang, O., Head-Gordon, T. & Forman-Kay, J. D. Biological insights from integrative modeling of intrinsically disordered protein systems. Curr Opin Struct Biol 93, 103063 (2025).

32. Bakker, M. J. et al. Streamlining NMR Chemical Shift Predictions for Intrinsically Disordered Proteins: Design of Ensembles with Dimensionality Reduction and Clustering. J Chem Inf Model 64, 6542–6556 (2024).

33. Tamiola, K., Acar, B. & Mulder, F. A. A. Sequence-specific random coil chemical shifts of intrinsically disordered proteins. J Am Chem Soc 132, 18000–18003 (2010).

34. Kjaergaard, M. & Poulsen, F. M. Disordered proteins studied by chemical shifts. Prog Nucl Magn Reson Spectrosc 60, 42–51 (2012).

35. Yao, J., Dyson, H. J. & Wright, P. E. Chemical shift dispersion and secondary structure prediction in unfolded and partly folded proteins. FEBS Lett 419, 285–289 (1997).

36. Tamiola, K., Acar, B. & Mulder, F. A. A. Sequence-specific random coil chemical shifts of intrinsically disordered proteins. J Am Chem Soc 132, 18000–18003 (2010).

37. Robustelli, P., Cavalli, A. & Vendruscolo, M. Determination of Protein Structures in the Solid State from NMR Chemical Shifts. Structure 16, 1764–1769 (2008).

38. Case, D. A. The use of chemical shifts and their anisotropies in biomolecular structure determination. Curr Opin Struct Biol 8, 624–630 (1998).

39. Cavalli, A., Salvatella, X., Dobson, C. M. & Vendruscolo, M. Protein structure determination from NMR chemical shifts. Proc Natl Acad Sci U S A 104, 9615–9620 (2007).

40. Robustelli, P., Kohlhoff, K., Cavalli, A. & Vendruscolo, M. Using NMR chemical shifts as structural restraints in molecular dynamics simulations of proteins. Structure 18, 923–933 (2010).

41. Camilloni, C., Cavalli, A. & Vendruscolo, M. Assessment of the use of NMR chemical shifts as replica-averaged structural restraints in molecular dynamics simulations to characterize the dynamics of proteins. Journal of Physical Chemistry B 117, 1838–1843 (2013).

42. Krieger, J. M. et al. Conformational recognition of an intrinsically disordered protein. Biophys J 106, 1771–1779 (2014).

43. Kleywegt, G. J. Validation of protein models from Cα coordinates alone. J Mol Biol 273, 371–376 (1997).

44. Tozzini, V., Rocchia, W. & McCammon, J. A. Mapping all-atom models onto one-bead coarse-grained models: General properties and applications to a minimal polypeptide model. J Chem Theory Comput 2, 667–673 (2006).

45. Kohlhoff, K. J., Robustelli, P., Cavalli, A., Salvatella, X. & Vendruscolo, M. Fast and accurate predictions of protein NMR chemical shifts from interatomic distances. J Am Chem Soc 131, 13894–13895 (2009).

46. Shen, Y. & Bax, A. SPARTA+: a modest improvement in empirical NMR chemical shift prediction by means of an artificial neural network. https://doi.org/10.1007/s10858-010-9433-9 doi:10.1007/s10858-010-9433-9.

47. Han, B., Liu, Y., Ginzinger, S. W. & Wishart, D. S. SHIFTX2: Significantly improved protein chemical shift prediction. J Biomol NMR 50, 43–57 (2011).

48. Li, J., Bennett, K. C., Liu, Y., Martin, M. V. & Head-Gordon, T. Accurate prediction of chemical shifts for aqueous protein structure on “Real World” data. Chem Sci 11, 3180–3191 (2020).

49. Qi, G. et al. Enhancing Biomolecular Simulations with Hybrid Potentials Incorporating NMR Data. J Chem Theory Comput 18, 7733–7750 (2022).

50. Cullen, M. et al. Integrating NMR restraints into coarse-grained simulations: toward accurate conformational ensembles of complex protein systems. bioRxiv 2025.12.22.695971 (2025) doi:10.64898/2025.12.22.695971.

51. Tesei, G., Schulze, T. K., Crehuet, R. & Lindorff-Larsen, K. Accurate model of liquid-liquid phase behavior of intrinsically disordered proteins from optimization of single-chain properties. Proc Natl Acad Sci U S A 118, e2111696118 (2021).

52. Montserret, R., McLeish, M. J., Böckmann, A., Geourjon, C. & Penin, F. Involvement of Electrostatic Interactions in the Mechanism of Peptide Folding Induced by Sodium Dodecyl Sulfate Binding†,‡. Biochemistry 39, 8362–8373 (2000).

53. Tesei, G. et al. Conformational ensembles of the human intrinsically disordered proteome. Nature 626, 897–904 (2024).

54. Sanz-Hernández, M. & De Simone, A. The PROSECCO server for chemical shift predictions in ordered and disordered proteins. J Biomol NMR 69, 147–156 (2017).

55. Gopinathan Nair, A. et al. Unorthodox PCNA Binding by Chromatin Assembly Factor 1. Int J Mol Sci 23, 11099 (2022).

56. Borysik, A. J., Kovacs, D., Guharoy, M. & Tompa, P. Ensemble Methods Enable a New Definition for the Solution to Gas-Phase Transfer of Intrinsically Disordered Proteins. J Am Chem Soc 137, 13807–13817 (2015).

57. Johansen, D., Trewhella, J. & Goldenberg, D. P. Fractal dimension of an intrinsically disordered protein: Small-angle X-ray scattering and computational study of the bacteriophage λ N protein. Protein Science 20, 1955–1970 (2011).

58. Launay, H. et al. Absence of residual structure in the intrinsically disordered regulatory protein CP12 in its reduced state. Biochem Biophys Res Commun 477, 20–26 (2016).

59. Gibbs, E. B. & Showalter, S. A. Quantification of Compactness and Local Order in the Ensemble of the Intrinsically Disordered Protein FCP1. Journal of Physical Chemistry B 120, 8960–8969 (2016).

60. Kiss, R., Kovács, D., Tompa, P. & Perczel, A. Local structural preferences of calpastatin, the intrinsically unstructured protein inhibitor of calpain. Biochemistry 47, 6936–6945 (2008).

61. Moffat, L. & Jones, D. T. Increasing the accuracy of single sequence prediction methods using a deep semi-supervised learning framework. Bioinformatics 37, 3744–3751 (2021).

62. Bjarnason, S. et al. DNA binding redistributes activation domain ensemble and accessibility in pioneer factor Sox2. Nat Commun 15, (2024).

63. Venters, R. A. et al. Use of 1HN- 1HN NOEs to Determine Protein Global Folds in Perdeuterated Proteins. J Am Chem Soc 117, 9592–9593 (1995).

64. Sanz-Hernández, M. & De Simone, A. The PROSECCO server for chemical shift predictions in ordered and disordered proteins. J Biomol NMR 69, 147 (2017).

65. Cao, F., von Bülow, S., Tesei, G. & Lindorff-Larsen, K. A coarse-grained model for disordered and multi-domain proteins. Protein Science 33, (2024).

66. Regy, R. M., Thompson, J., Kim, Y. C. & Mittal, J. Improved coarse-grained model for studying sequence dependent phase separation of disordered proteins. Protein Science 30, 1371–1379 (2021).

67. Latham, A. P. & Zhang, B. Consistent Force Field Captures Homologue-Resolved HP1 Phase Separation. J Chem Theory Comput 17, 3134–3144 (2021).

68. Holehouse, A. S. & Kragelund, B. B. The molecular basis for cellular function of intrinsically disordered protein regions. Nature Reviews Molecular Cell Biology 2023 25:3 25, 187–211 (2023).

69. Ruff, K. M. et al. Molecular grammars of predicted intrinsically disordered regions that span the human proteome. Cell 0, (2025).

70. Ginell, G. M. et al. Sequence-based prediction of intermolecular interactions driven by disordered regions. Science (1979) 388, (2025).

71. Wang, X., Ramírez-Hinestrosa, S., Dobnikar, J. & Frenkel, D. The Lennard-Jones potential: when (not) to use it. Physical Chemistry Chemical Physics 22, 10624–10633 (2020).

72. Novak, B. et al. Accurate predictions of conformational ensembles of disordered proteins with STARLING. bioRxiv 2025.02.14.638373 (2025) doi:10.1101/2025.02.14.638373.

73. Tesei, G., Pesce, F. & Lindorff-Larsen, K. Computational design of intrinsically disordered proteins. https://arxiv.org/abs/2509.12460v1 (2025).

74. Liu, G. et al. NMR data collection and analysis protocol for high-throughput protein structure determination. Proc Natl Acad Sci U S A 102, 10487 (2005).

75. Baskaran, K. et al. Restraint validation of biomolecular structures determined by NMR in the Protein Data Bank. Structure 32, 824–837.e1 (2024).

76. Ulrich, E. L. et al. BioMagResBank. Nucleic Acids Res 36, D402–D408 (2008).

77. De Simone, A., Cavalli, A., Danny Hsu, S.-T., Vranken, W. & Vendruscolo, M. Accurate Random Coil Chemical Shifts from an Analysis of Loop Regions in Native States of Proteins. https://doi.org/10.1021/ja904937a doi:10.1021/ja904937a.

78. Clevert, D. A., Unterthiner, T. & Hochreiter, S. Fast and Accurate Deep Network Learning by Exponential Linear Units (ELUs). 4th International Conference on Learning Representations, ICLR 2016 - Conference Track Proceedings https://arxiv.org/abs/1511.07289v5 (2015).

79. Eastman, P. et al. OpenMM 8: Molecular Dynamics Simulation with Machine Learning Potentials. Journal of Physical Chemistry B 128, 109–116 (2024).

80. Bulacu, M. et al. Improved angle potentials for coarse-grained molecular dynamics simulations. J Chem Theory Comput 9, 3282–3292 (2013).

81. Torda, A. E. & van Gunsteren, W. F. The refinement of NMR structures by molecular dynamics simulation. Comput Phys Commun 62, 289–296 (1991).

82. Torda, A. E., Scheek, R. M. & van Gunsteren, W. F. Time-dependent distance restraints in molecular dynamics simulations. Chem Phys Lett 157, 289–294 (1989).

83. Marsh, J. A., Singh, V. K., Jia, Z. & Forman-Kay, J. D. Sensitivity of secondary structure propensities to sequence differences between α- and γ-synuclein: Implications for fibrillation. Protein Science 15, 2795–2804 (2006).

84. Dima, R. I. & Thirumalai, D. Asymmetry in the Shapes of Folded and Denatured States of Proteins†. Journal of Physical Chemistry B 108, 6564–6570 (2004).

85. Galvanetto, N. et al. Extreme dynamics in a biomolecular condensate. Nature 2023 619:7971 619, 876–883 (2023).

86. Humphrey, W., Dalke, A. & Schulten, K. VMD: Visual molecular dynamics. J Mol Graph 14, 33–38 (1996).

87. Meng, E. C. et al. UCSF ChimeraX: Tools for structure building and analysis. Protein Science 32, e4792 (2023).

