## SupplementaryInformation for "Capturing secondary structure in coarse grained intrinsically disordered proteins with simulations driven by chemical shifts"

### This document contains:

Figures S1 to S8.

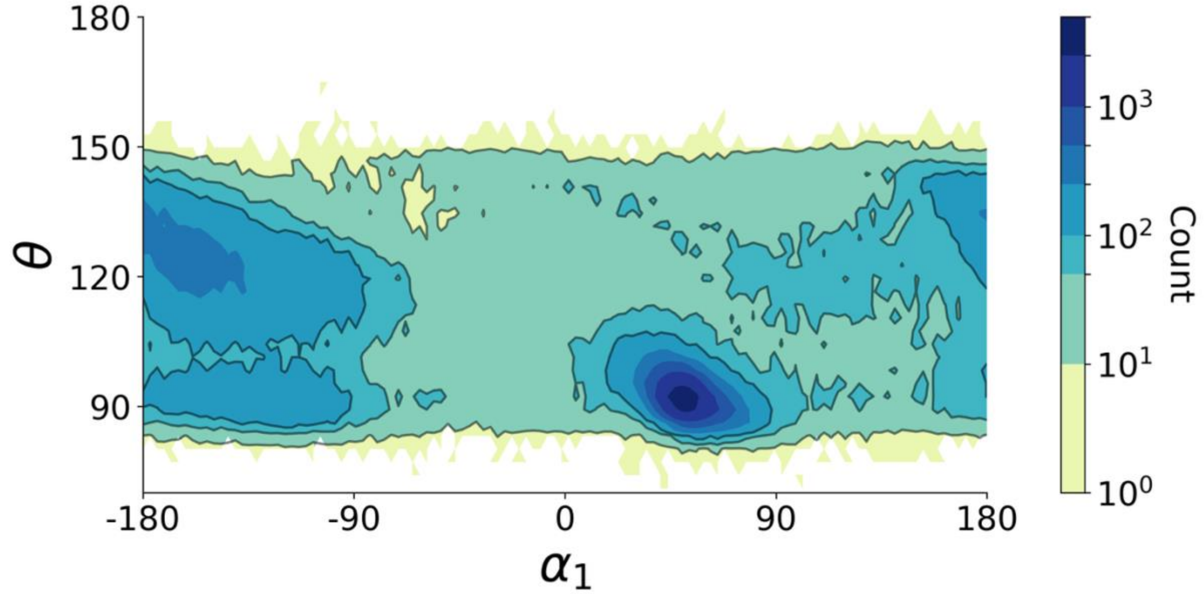


**Supplementary Figure 1 | Pseudoramachandran plot of NapshiftCα training set.** Pseudoramachandran (c) plot of θ vs. $\alpha_1$ of the dataset used to train Napshift$C_\alpha$, with entries coloured by count. The respective θ vs. $\alpha_2$ plot is reported in Figure 1c.

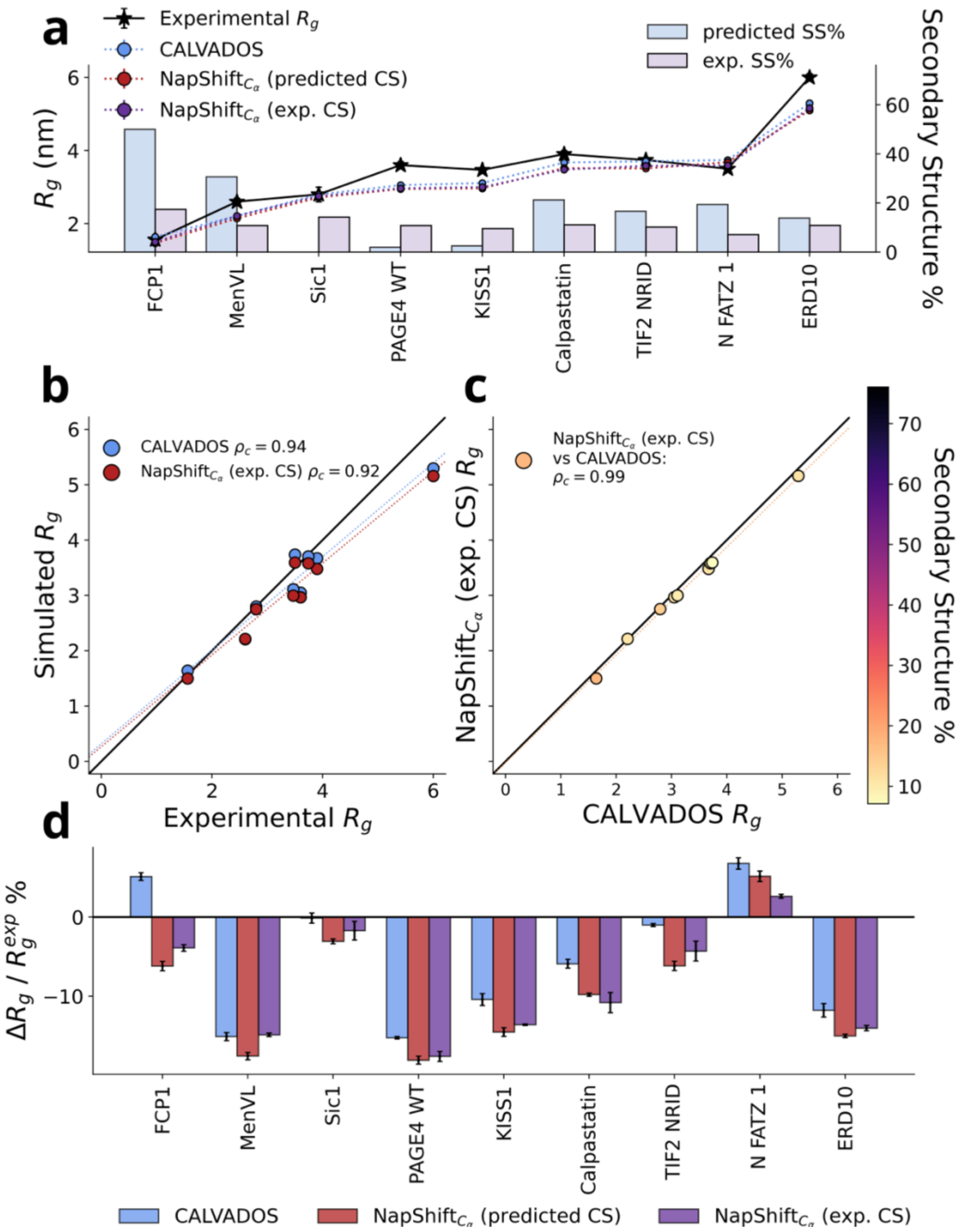


**Supplementary Figure 2 | Validation of NapshiftCα via experimental chemical shifts.** a) Experimental (stars) and simulated (dots) $R_g$ for proteins simulated using experimental CS (red) unrestrained simulations are shown as blue dots. The bars show the percentage of secondary structure obtained from CS predictions made with S4pred[1] (light blue) or obtained from experimental CS (violet). (b-c) concordance correlation coefficient between simulated and experimental Rg (b) and between restrained and unrestrained simulations (c). (d) Agreement between experimental and computed Rg as a function of the simulated proteins.

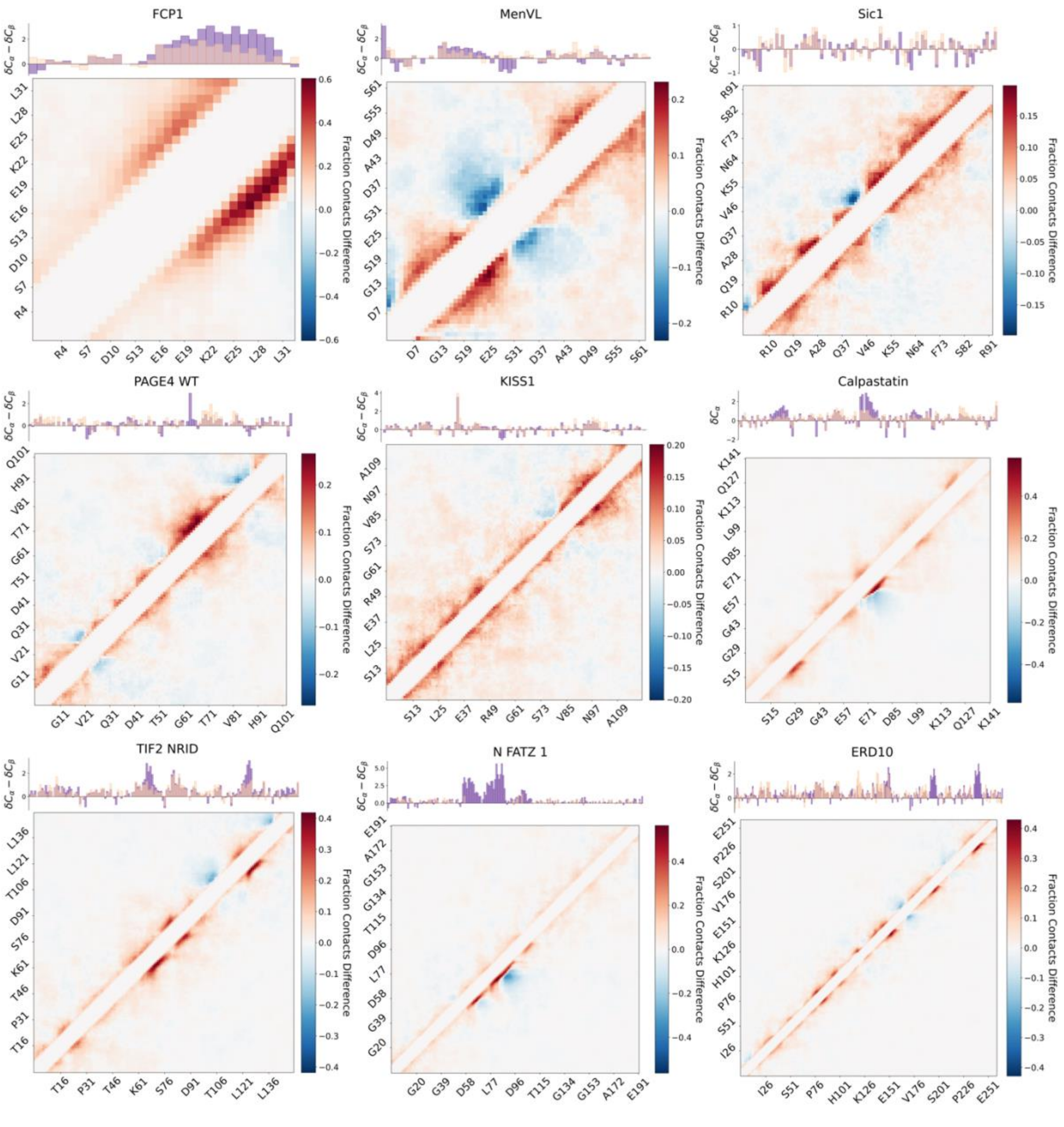


**Supplementary Figure 3 | Contact maps of ensembles simulated with experimental CS.** Each contact map is reported as the difference in the fraction of contacts obtained from subtracting unrestrained from restrained simulations. Blue and red regions show Napshift$C_\alpha$ making more and less contacts compared to Calvados, respectively. On the top of each map average experimental (pink) and predicted (purple) secondary chemical shifts are shown.

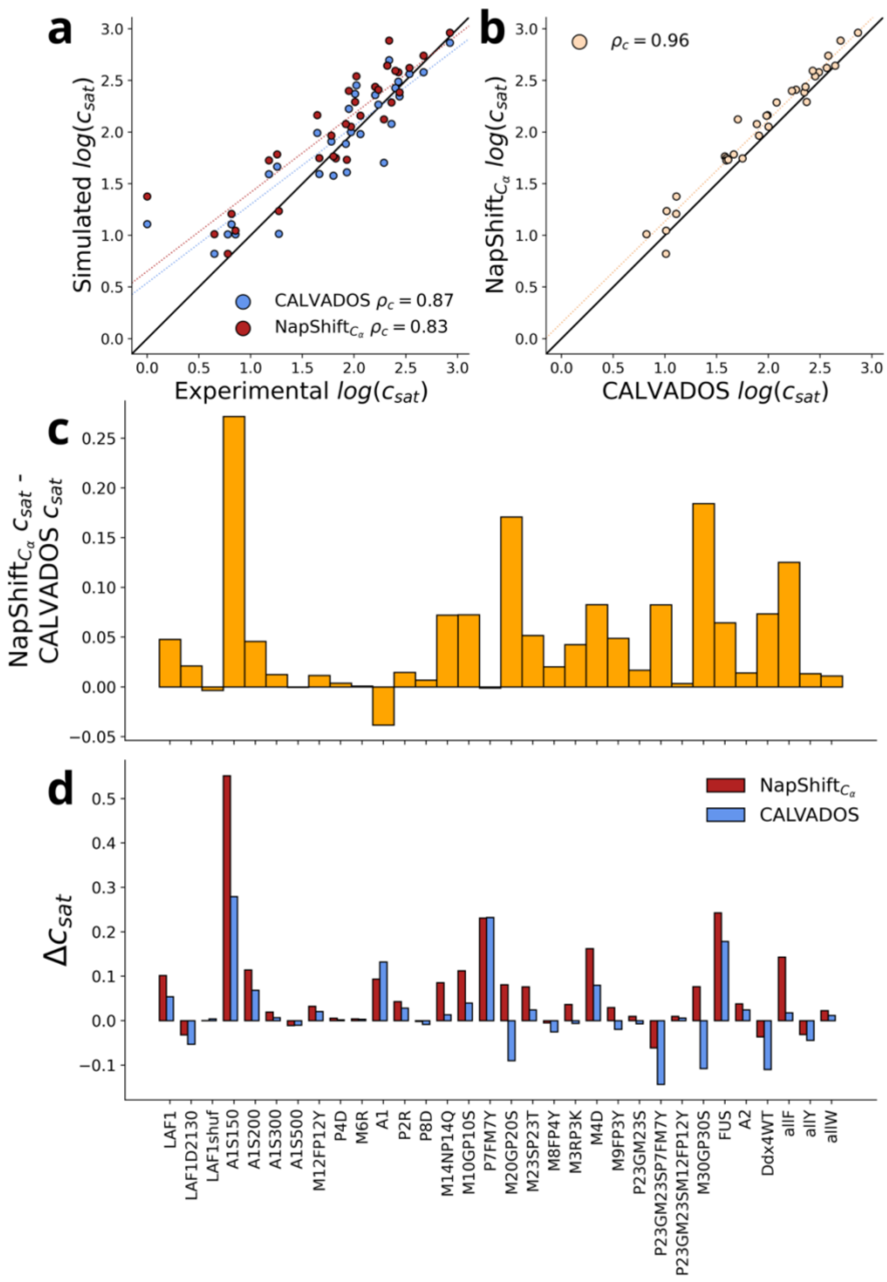


**Supplementary Figure 4 | Agreement of simulations with concentration of saturation ($c_{sat}$).** (a) correlation and concordance correlation coefficients between simulated and experimental values of $c_{sat}$ (a) and between restrained and unrestrained simulations (b). The dashed lines report a linear fit of the data. (c) $c_{sat}$ difference between restrained and unrestrained simulations per simulated protein. (d) difference ($\Delta c_{sat}$) between the $c_{sat}$ obtained from subtracting restrained (red) from unrestrained (blue) simulations and experiments, per simulated protein.

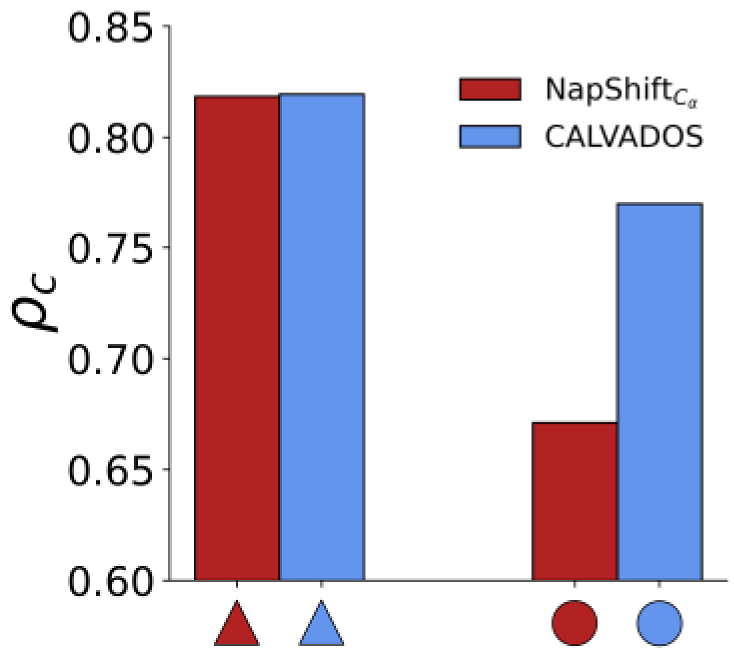


**Supplementary Figure 5 | Agreement between experimental and computed FRET efficiencies for regions of eIF4b encompassing secondary structure or disordered.** Concordance correlation coefficients ($\rho_c$) for the computed and experimental <E> for the regions of eIF4b either encompassing secondary structure (triangles) or falling at the C-terminal end (random coil) of the protein (dots).

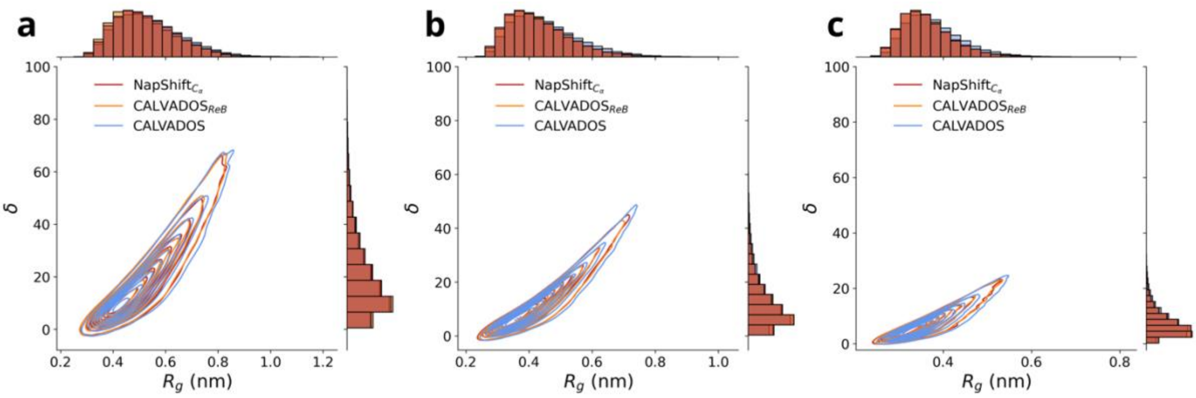


**Supplementary Figure 6 | Overall dimensions of simulated proteins.** Asphericity (δ) as a function of the $R_g$ for Sox2 (a), the Protα-Histone H1 (b) and eIF4b (c), obtained from simulations as reported in the legend.

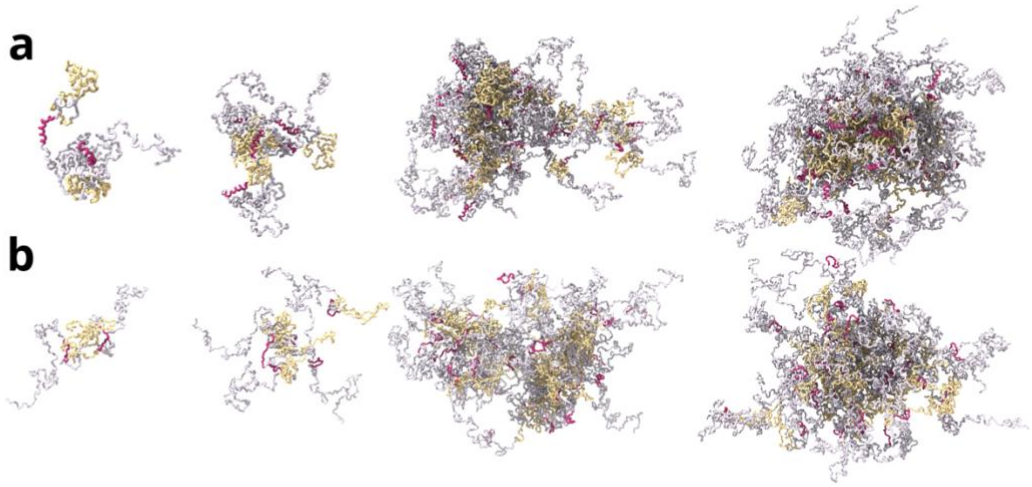


**Supplementary Figure 7 | Ensemble of condensates formed by eIF4b.** (a) NapshiftCα-restrained Calvados simulations. (b) Unrestrained Calvados simulations. The DRYG domain is shown in yellow, the region of the molecule corresponding to the α-helical segment is shown in red, while the rest of the chain is shown in grey.

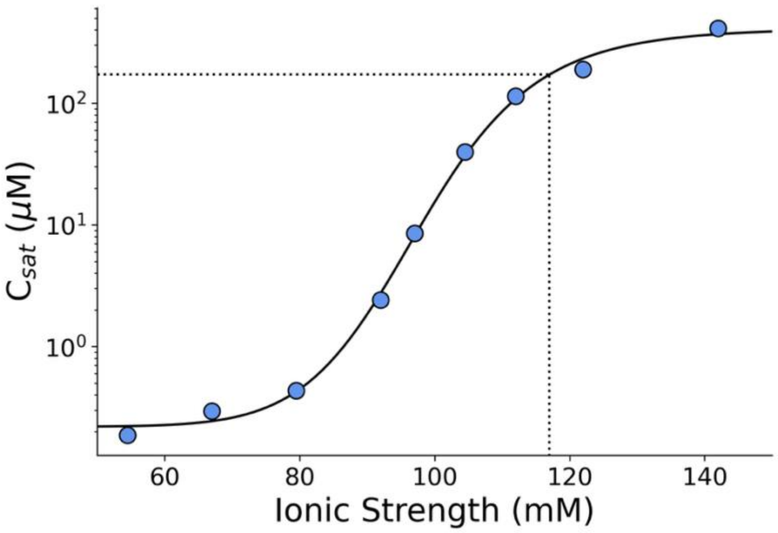


**Supplementary Figure 8 | concentration of saturation as a function on ionic strength in eIF4b condensates.** The dashed line shows the value at which simulations shown in Figure 5 were performed.

### Supplementary information references

1. Moffat, L. & Jones, D. T. Increasing the accuracy of single sequence prediction methods using a deep semi-supervised learning framework. *Bioinformatics* 37, 3744–3751 (2021).
